# *In vitro* synergy between Sodium Deoxycholate and Furazolidone against Enterobacteria via Inhibition of Multidrug Efflux Pumps

**DOI:** 10.1101/518050

**Authors:** Vuong Van Hung Le, Catrina Olivera, Julian Spagnuolo, Ieuan Davies, Jasna Rakonjac

**Affiliations:** School of Fundamental Sciences, Massey University, Palmerton North, New Zealand; New Zealand Pharmaceutical Ltd., Palmerston North, New Zealand

## Abstract

Antimicrobial combinations have been proven to be a promising approach in the confrontation with multi-drug resistant bacterial pathogens, owing to enhancement of antibacterial efficacy, deceleration of resistance development rate and mitigation of side effects by lowering the doses of two drugs. In the present study, we report that combination of furazolidone (FZ) and other nitrofurans with a secondary bile salt, Sodium Deoxycholate (DOC), generates a profound synergistic effect on growth inhibition and lethality in enterobacteria, including *Escherichia coli*, *Salmonella*, *Citrobacter gillenii* and *Klebsiella pneumoniae*. Taking *E. coli* as the model organism to study the mechanism of DOC-FZ synergy, we found that the synergistic effect involves FZ-mediated inhibition of efflux pumps that normally remove DOC from bacterial cells. We further show that the FZ–mediated nitric oxide production contributes to the synergistic effect. This is to our knowledge the first report of nitrofuran-DOC synergy against Gram-negative bacteria.

## Introduction

Antimicrobial resistance (AMR) is one of the most serious threats with which humans have been confronted. A UK-Prime-Minister-commissioned report in 2014 estimated that AMR, without appropriate interventions, will cause globally 10 million deaths per annum with a cumulative loss of US $100 trillion by 2050 (1). In this dire context, alternative approaches are urgently needed besides traditional discovery of novel antibiotics, in which antimicrobial combinations have been proven to be a promising approach with some widely accepted advantages, including enhancement of antimicrobial efficacy, deceleration of the rate of resistance and alleviation of side effects (2, 3). Moreover, this approach could amplify the significance of ongoing antimicrobial discovery programs; particularly the advent of any novel antimicrobial compound would bring about a large number of possible double combinations with existing antimicrobial agents to be evaluated, let alone triple and quadruple combinations.

Sodium Deoxycholate (DOC) (Figure 1E) is a facial amphipathic compound in bile, which is secreted into the duodenum to aid lipid digestion and confer some antimicrobial protection (4). Though extensive research has been conducted to elucidate the interaction between DOC, either alone or in bile mixture, and enteric bacteria, the mode of its antimicrobial action remains elusive. It was suggested that DOC could attack multiple cellular targets, including disturbing cell membranes, causing DNA damage, triggering oxidative stress and inducing protein misfolding (4–6). Nonetheless, Gram-negative bacteria such as *Escherichia coli* and *Salmonella* are highly resistant to DOC by many mechanisms such as employment of diverse active efflux pumps, down-regulation of outer membrane porins and activation of various stress responses (5, 7-10).

**Figure 1:**
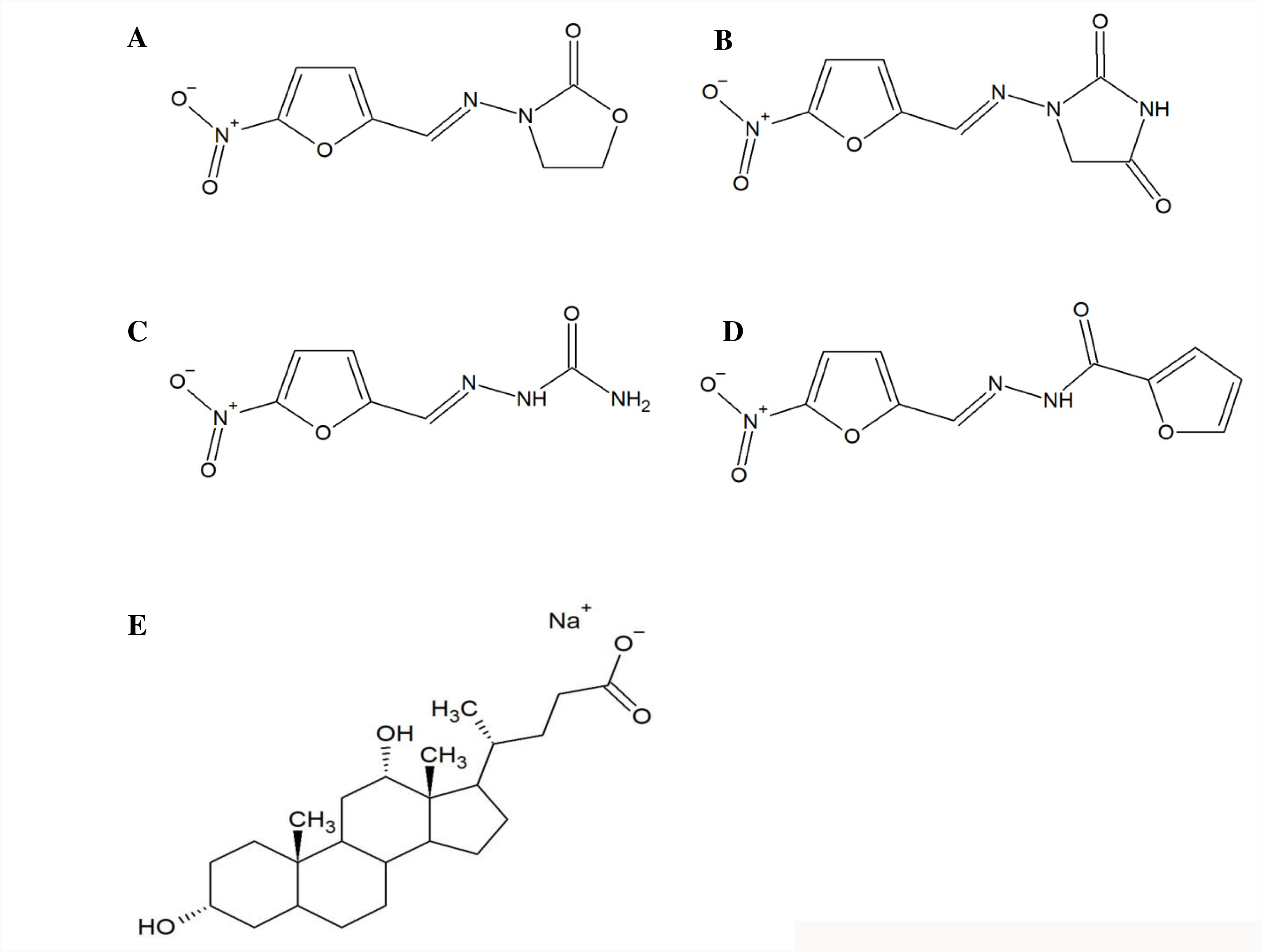
Structural formulae of nitrofurans and Sodium Deoxycholate (E). A) Furazolidone (FZ); B) Nitrofurantoin (NIT); C) Nitrofurazone (NFZ). D) CM4, Pubchem ID AC1LGLMG (no CAS number). Chemical name: N'-[(5-nitrofuran-2-yl)methylidene]furan-2-carbohydrazide or N-[(5-nitrofuran-2-yl)methylideneamino]furan-2-carboxamide.

The 5-nitrofurans are an old class of synthetic antimicrobials, clinically introduced in the 1940s and 1950s (11); several are commercially available, including furazolidone (FZ), nitrofurantoin (NIT) and nitrofurazone (NFZ) (Figure 1). FZ is used to treat bacterial diarrhea, giardiasis and as a component in combinatorial therapy for *Helicobacter pylori* infections; NIT and NFZ are used for urinary tract infections and topical applications, respectively (12). They are prodrugs which require reductive activation mediated largely by two type-I oxygen-insensitive nitroreductases, NfsA and NfsB. These two enzymes perform stepwise 2-electron reduction of the nitro moiety of the compound into the nitroso and hydroxylamino intermediates and biologically inactive amino-substituted product (13, 14). The detailed mechanism of how bacterial cells are killed by the reactive intermediate has yet to be clarified. Nevertheless, it has been proposed that the hydroxylamino derivatives could trigger DNA lesions, disrupt protein structure and arrest RNA and protein biosynthesis (15-19). Some reports also suggested that nitric oxide could be generated during the activation process, thus inhibiting electron transport chain of bacterial cells though clear evidence for that is not available as yet (20, 21).

In this study, we have characterized interaction of DOC with FZ and three other related nitrofurans against a range of enterobacteria. We identified the underlying mechanism of DOC-FZ synergy using *E. coli* K12 as a model organism.

## Results

### The synergy between DOC and 5-nitrofurans against enterobacteria

To evaluate the synergy between DOC and FZ, the checkerboard growth inhibition assays were performed for a range of enterobacteria, including *Salmonella enterica* sv. Typhimurium LT2, *Citrobacter gillenii, Klebsiella pneumoniae* and two *E. coli* antibiotic-resistant strains (streptomycin-resistant and streptomycin/ampicillin-resistant). DOC and FZ act synergistically in inhibiting the growth of the microorganisms listed (Figure 2), with FICI ranging from 0.125 in streptomycin-resistant *E. coli* strain (Figure 2**A**) to 0.35 in *K. pneumoniae* (Figure 2F). DOC-FZ synergy was also observed against two *E. coli* pathogenic strains (*E. coli* strain O157 and urinary tract infection strain P50; Figure S1). It is worth noting that, when used alone, very high DOC concentrations were required to exert an equivalent effect on inhibiting the growth of these Gram-negative enterobacteria, reflecting the inherent resistance to DOC in these bacteria thanks to their impermeable outer membrane and active efflux pumps, which prevent the intracellular accumulation of toxic xenobiotics.

**Figure 2:**
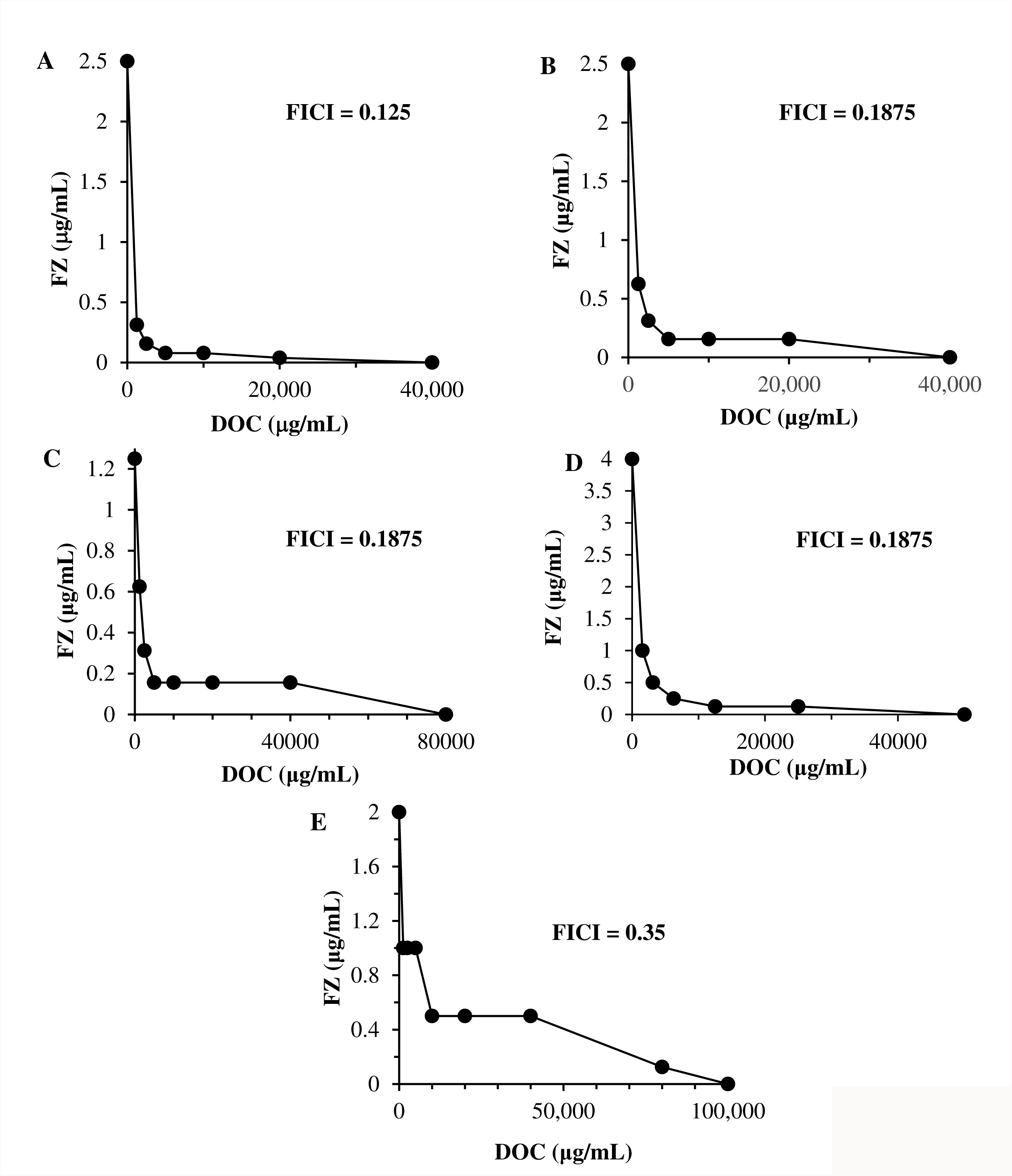
FZ interaction with DOC in growth inhibition of streptomycin-resistant *E. coli* K12 (A), ampicillin- and streptomycin-resistant *E. coli* K12 (B), *Salmonella enterica* sv. Typhimurium LT2 (C), *Citrobacter gillenii* (D) and *Klebsiella pneumoniae* (E). Graphs (isobolograms) are obtained using a checkerboard analysis at multiple concentration of molecules. Each data point represents the minimum molecule concentrations alone or in combination causing 90 % inhibition to bacterial growth.

We also examined the interaction between DOC and other nitrofuran compounds, including NIT, NFZ and CM4 (a 5-nitrofuran compound we found during an antimicrobial synergy screening campaign against *E. coli*, Figure 1D) in all the bacterial species mentioned above. We found that NIT, NFZ and CM4 were synergistic with DOC in *E. coli* laboratory strain (Figure 4), *Citrobacter gillenii* (Figure S2) and *Salmonella* Typhimurium LT2 (Figure S3) but indifferent in *K. pneumoniae* isolate (Figure S4).

**Figure 4:**
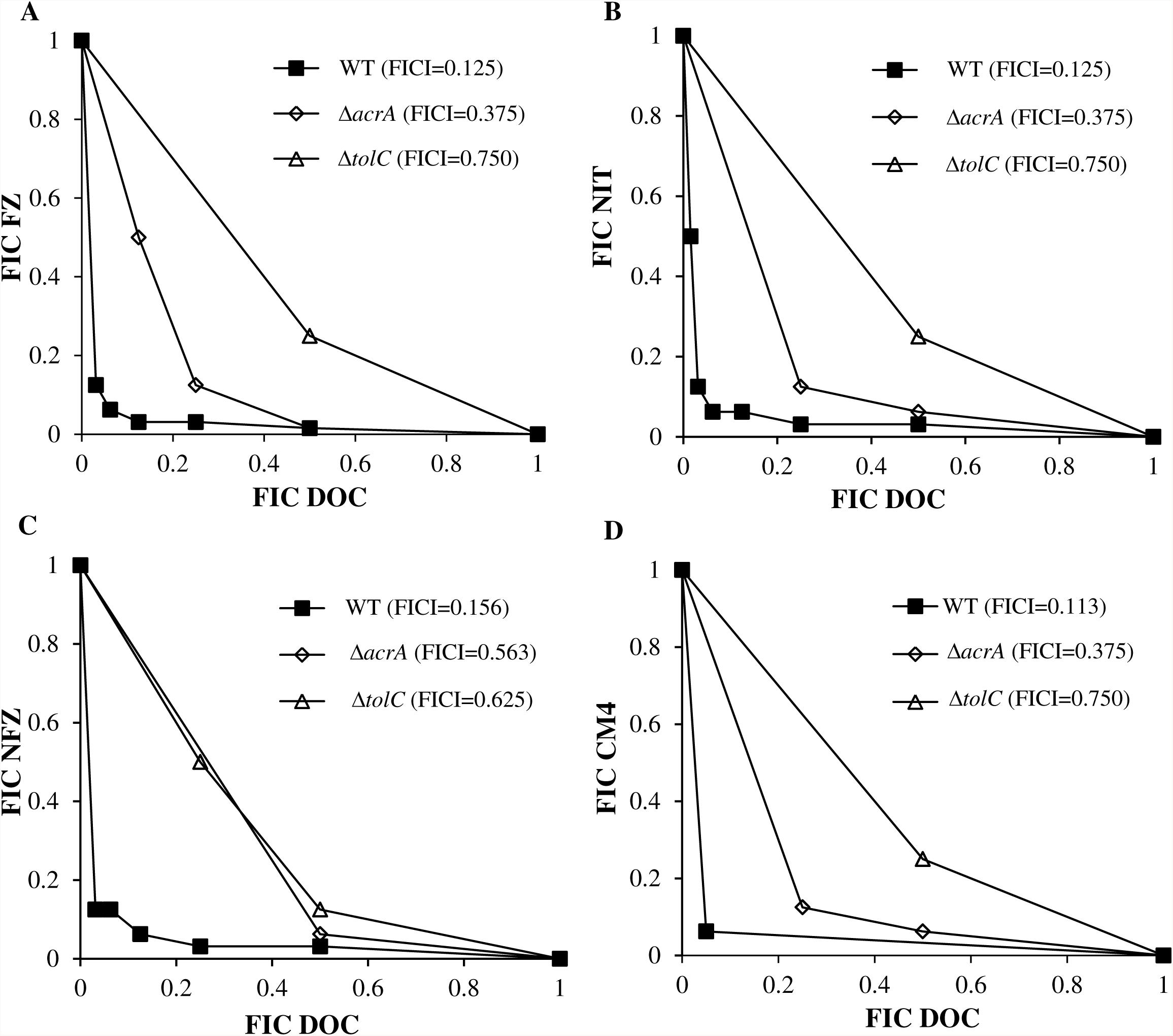
Effect of the Δ*tolC* and *ΔacrA* mutations on DOC synergy with FZ, NIT, NFZ and CM4 in *E. coli*. Isobolograms characterizing interactions of DOC with FZ (A), NIT (B), NFZ (C) and CM4 (D) in growth inhibition assays of the *E. coli* K12 strain K1508 (WT or wild-type and two isogenic deletion mutants, Δ*acrA* and Δ*tolC*). Each data point corresponds to the FIC (ratios of the 90% growth inhibition concentrations in combination vs. alone) for one of the four nitrofurans (y axis) and DOC (x axis).

To elaborate the interaction between DOC and FZ in terms of bactericidal effects, the time-kill assay was employed. Streptomycin-resistant *E. coli* K12 laboratory strain K1508 and *S. enterica* serovar Typhimurium strain LT2 were exposed to sub-inhibitory concentrations of DOC (2500 µg/mL) alone, or FZ (0.5 × MIC) alone, or combination of the two drugs at such sub-inhibitory concentrations, over a 24 h period. The sample was taken at different time points and the surviving bacteria were titrated on the antimicrobial-free plates. Centrifugation and resuspension were applied for each sample to eliminate the antimicrobial carryover before plating. After 24 h, the total cell count in the sample treated with the DOC-FZ combination was about six to seven orders of magnitude lower than that in the sample treated with either DOC or FZ alone for both *E. coli* and *Salmonella* (Figure 3), demonstrating the synergy in bacterial killing between DOC and FZ.

**Figure 3:**
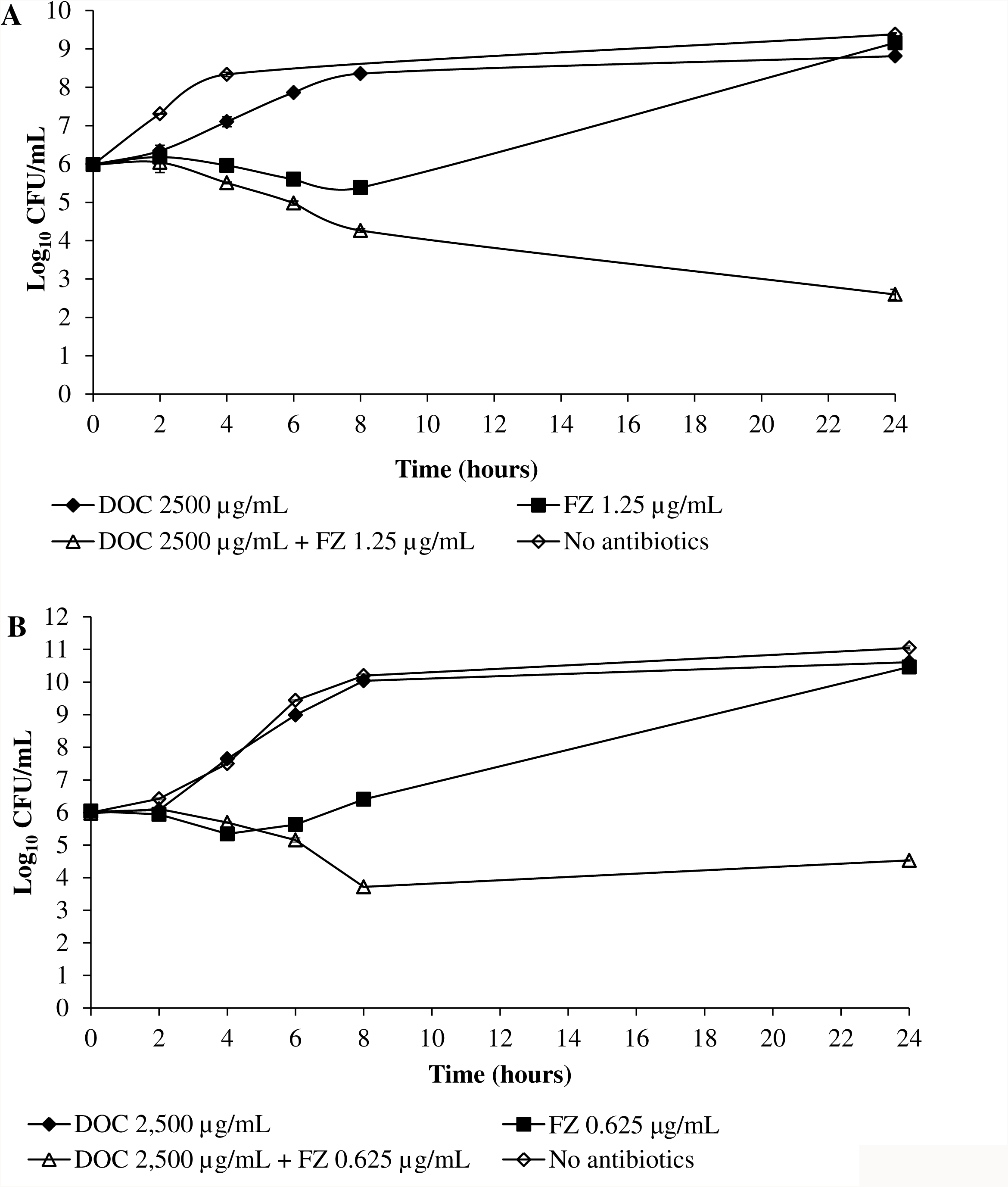
Time-kill analysis of the DOC and FZ combination in killing *E. coli* strain K1508 (A) and *Salmonella enterica* sv. Typhimurium LT2 (B). The data is presented as the mean ± standard error of the mean (SEM) of three independent measurements. The count of the live cells was determined at indicated time points by titration of colony-forming units on agar plates. The lower limit of detection was 60 CFU/mL.

### The role of AcrAB-TolC efflux pump in synergistic interaction between DOC and nitrofurans

One commonly accepted principle is that the synergy between two drugs is a consequence of one drug suppressing bacterial physiological pathways that mediate resistance to the other one. It has been reported that DOC could be expelled out of the cell via a wide range of efflux pumps, in which the tripartite efflux system AcrAB-TolC plays the major role (7, 9). This led us to hypothesize that FZ inhibits the activity of efflux pumps, thus allowing intracellular accumulation of DOC to exert its lethal effect. If this scenario were true, disruption of the function of efflux pumps by mutation was expected to make this activity of FZ redundant, thus reducing the interaction index (FICI) in the mutant strains.

To validate this model in *E. coli*, the checkerboard assay was performed on the strains containing deletions of individual genes encoding the AcrAB-TolC efflux pump system, Δ*tolC* and Δ*acrA*. Deletion of *tolC* caused a shift from the synergistic interaction between DOC and FZ in the wild type (FICI = 0.125) to indifferent interaction (FICI=0.75; Figure 4A). The Δ*acrA* mutant exhibited a 3-fold decrease in the FICI index relative to the isogenic wild type strain. Such changes were also observed for the interaction between DOC and other nitrofurans, NIT, NFZ or CM4 (Figure 4BCD).

To confirm that these observations were conferred by direct effect of the *tolC* and *acrA* deletion, rather than indirect effects of other genes or proteins, complementation of the corresponding deletion mutations by plasmid-expressed *tolC* and *acrA* was performed. To compensate for the multiple copies of plasmid-containing genes, complementation was carried out at a low level of expression, nevertheless it completely restored the strong synergy between DOC and FZ in these complemented strains (Figure 5). These findings collectively support the model that the efflux pumps act as the interacting point for the synergy between DOC and FZ.

**Figure 5:**
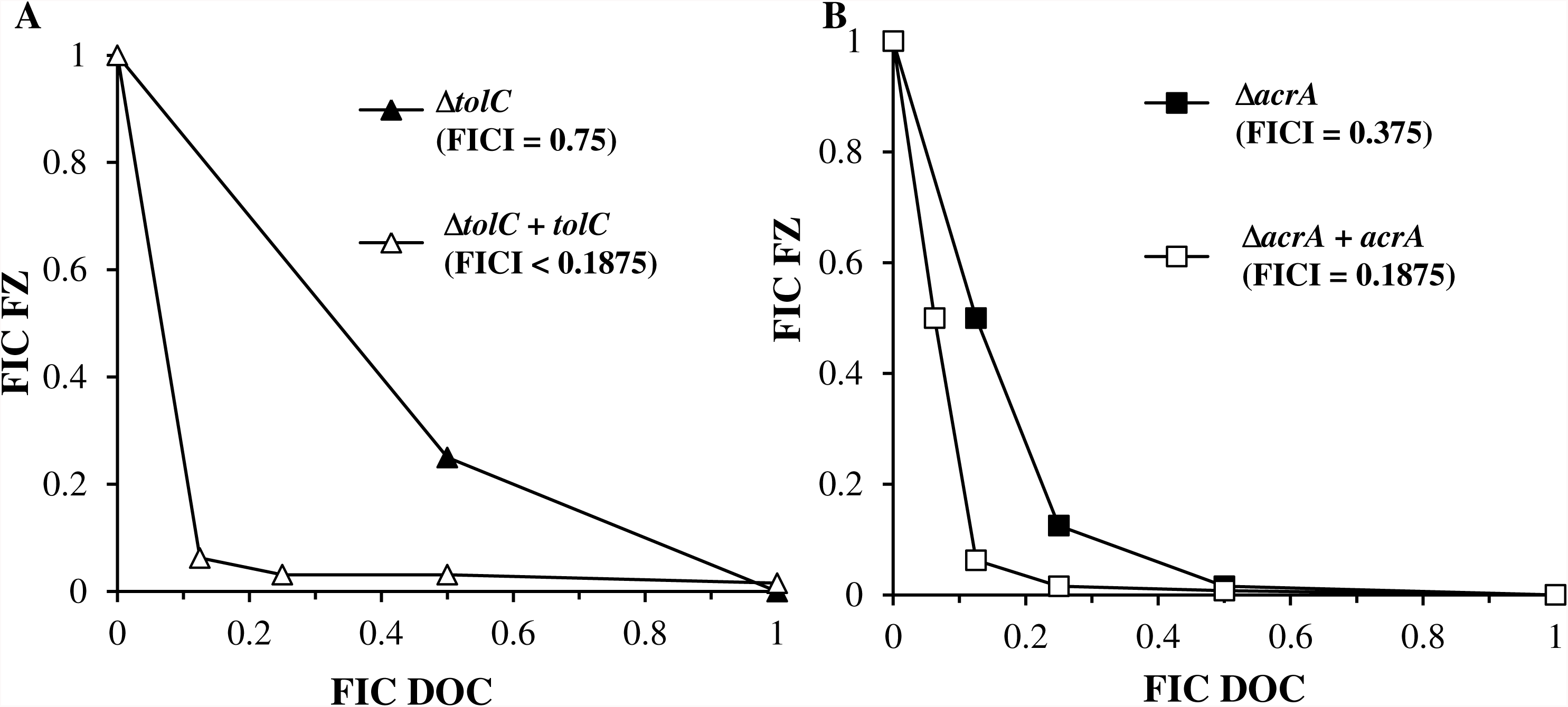
Recovery of FZ-DOC synergy in complemented Δ*tolC* and Δ*acrA* mutants. Isobolograms of FZ-DOC interactions in growth inhibition of: A. Δ*tolC* mutant (Δ*tolC*) and a derived strain containing a plasmid expressing *tolC* gene (Δ*tolC* + *tolC*); B. Δ*acrA* mutant (Δ*acrA*) and a derived strain containing a plasmid expressing *acrA* gene and (Δ*acrA* + *acrA*). Each data point corresponds to the FIC (ratios of the 90% growth inhibition concentrations in combination vs. alone) for FZ (y axis) and DOC (x axis).

An intriguing question to be unraveled is how FZ could negatively influence the action of efflux pumps. We hypothesized that FZ could lower the energy supply to efflux pumps by mediating an increase in concentration of nitric oxide (NO). To verify the proposed model, the interaction between DOC and FZ in the *E. coli* strain with increased expression of protein Hmp (the *E. coli* nitric oxide dioxygenase) was inspected. The rationale for this is that overexpression of Hmp protein would increase detoxification of NO by conversion into benign NO_3_^−^ions, thus relieving the effect exerted by NO (22). If NO was involved in the mechanism of the interaction between the two drugs, the synergy degree between them was expected to decrease with an increased abundance of Hmp proteins. In agreement with this hypothesis, overexpression of *hmp* was found to suppress the synergy between DOC and FZ by a factor of 3 (Figure 6). This finding supports the model that NO generated during FZ metabolism participates in the inhibition of electron transport chain (23), with the secondary effect of inhibiting the function of efflux pumps which are dependent on the electron transport chain for their activity.

**Figure 6:**
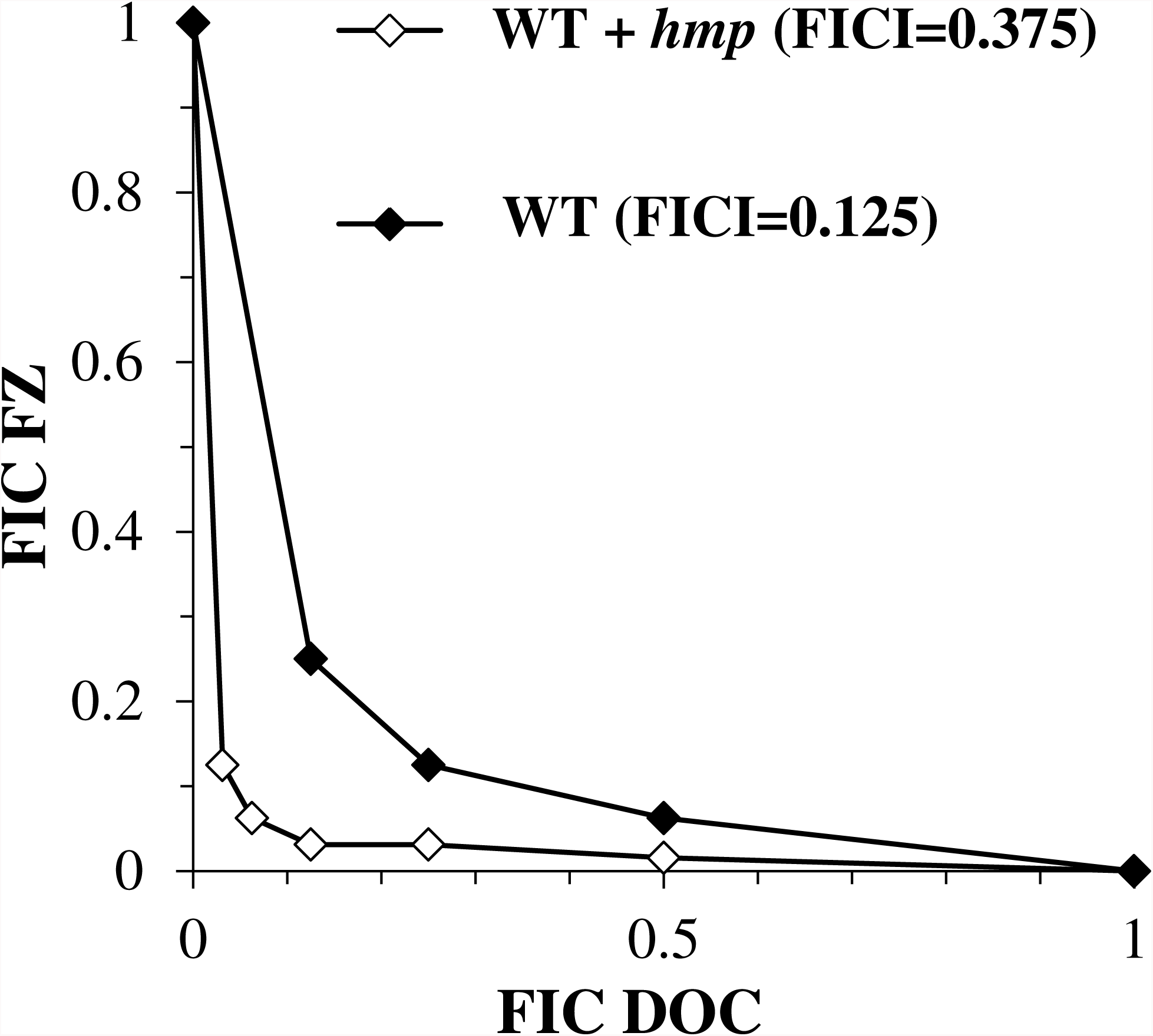
Effect of the *hmp* gene overexpression on FZ-DOC synergy. The isobologram of DOC and FZ interaction in *E. coli* having differential expression of NO-detoxifying protein Hmp. WT, strain *E. coli* laboratory strain K1508; WT + *hmp*, K1508 containing a plasmid expressing Hmp under the control of a T5-lac hybrid promoter. Expression of *hmp* gene was induced by IPTG (1 mM). Each data point corresponds to the FIC (ratios of the 90% growth inhibition concentrations in combination vs. alone) for FZ (y axis) and DOC (x axis).

## Discussion

The widespread emergence of antimicrobial drug resistance and the drying pipeline of antibiotics for Gram-negative pathogens imposed an urgency to seek for novel approaches to combat these pathogens. Capitalization on drug combinations is one of the promising approaches to design novel therapies that will allow application of antimicrobials which have heretofore been ineffective against Gram-negative bacteria at concentrations that are acceptable for medical treatments. In the present study, we describe the synergistic interaction between DOC and FZ in a range of enterobacteria in terms of growth inhibition and/or lethality. These findings offer two major implications. Firstly, Gram-negative bacteria, such as *E. coli* and *Salmonella* have evolved to be highly resistant to bile salts, including DOC (10); inclusion of an active agent, such as FZ or other 5-nitrofurans could revitalize DOC in the battle against such formidable pathogens. This discovery raises a possibility of using synergistic combinations in enabling use of antimicrobials that are on their own ineffective against Gram-negative bacteria at sub-toxic concentrations, for treatment of infections caused by these resilient organisms.

Secondly, DOC and other bile salts are inherently present at varying concentrations along the gastrointestinal tract. The efficacy of any drug dedicated to treat intestinal infections would depend on physicochemical properties of the local environment in which interaction with bile salts is one important factor. The synergy between DOC and FZ described here partly explains the success of using FZ in curing bacterial diarrhea (12, 24). To further highlight such an interaction, it has also been reported that rifaximin, an RNA synthesis inhibitor, worked more efficiently in treating diarrhea-producing *E. coli* in the intestine than in the colon, due to the difference in the bile salt concentration (25). From these observations, we propose the co-administration of DOC and FZ to treat bacterial diarrhea for the patients who have low intestinal concentrations of DOC due to malnourishment, disorders in enterohepatic circulation or intestinal absorption (4). Nonetheless, further investigations are required to justify the validity of that proposal.

In this study, we have also provided some insights into the underlying mechanism of the synergy between DOC and FZ in their antibacterial action against *E. coli* as a model Gram-negative bacterium. We showed that disruption of *tolC* or *acrA* gene caused a considerable decrease in the synergy between DOC and FZ in the corresponding mutants. The TolC protein, whose removal disrupts the synergy more strikingly, appears to be the key determinant of synergy.

The observed difference in the susceptibility to DOC/FZ combination between Δ*tolC* and Δ*acrA* mutants is in agreement with the fact that the TolC protein is shared by at least seven multidrug efflux pumps, while AcrA protein acts as the periplasmic connecting bridge for only two (26). Thus, deletion of *tolC* gene is expected to give rise to a more pronounced effect on the loss of efflux activities than deletion of *acrA* gene.

Of great interest is how FZ could influence the activity of efflux pumps. The obtained findings indicate that more than two efflux pumps (AcrAB-TolC and AcrAD-TolC systems) were affected by FZ. This observation is reminiscent of a common mechanism which could affect a wide range of efflux pumps simultaneously, namely proton motive force. It has been suggested that nitrofuran compounds during reductive activation might generate nitric oxide (NO) which subsequently inhibits the electron transport chain (ETC), diminishing the proton motive force across the cytoplasmic membrane (20, 21, 23, 27). As a result, many efflux pumps would be de-energized, and become less efficient in extruding toxic compounds.

However, NO generation from nitrofurans in bacterial cells remains to be speculative since the trace of NO has yet to be detected using either biochemical or NO-sensing fluorescence methods, possibly due to the detection limit of the used methods or rapid conversion of NO into other compounds (20, 21). In the present work, we provide evidence for the contribution of NO in the interaction between DOC and FZ via the observation that overexpression of NO-detoxifying enzyme Hmp decreased the synergistic interaction between the two agents. Since some DOC-FZ synergy was still retained after NO-detoxification, other mechanisms, including direct inhibition of the ETC by activated FZ, are involved in the efflux pump inhibition.

In conclusion, we report the synergy between FZ and DOC in inhibiting and/or killing different enterobacterial species. In the terms of underlying mechanisms, much evidence supports the model that FZ negatively influences the activity of many efflux pumps such that DOC could accumulate inside the cell to exert its cytotoxic effect. One possible route is via FZ-derived NO which inhibits the electron transport chain, thus dissipating the energy supply for efflux machineries. Nonetheless, other mechanisms might be involved, remaining to be elucidated.

## Materials and methods

### Bacterial strains, growth conditions and antibiotics

All bacterial strains and plasmids used in this study were described in **Table 1** and **Table 2**. The introduction of the *kan^R^* gene deletion mutations into the wild type strain K1508 from the corresponding Keio collection *E. coli* K12 knock-out strains (28) was performed using phage P1 transduction, according to the standard procedures (29). To eliminate potential polar effects on downstream genes in the operon, the FRT-flanked *kan^R^* cassette was then removed using FLP-mediated recombination as previously described (30). Plasmids derived from the pCA24N bearing the gene of interest were purified from *E. coli* strains of the ASKA collection containing ORF expression constructs derived from this organism (31) using the ChargeSwitch-Pro Plasmid Miniprep Kit (Thermo Fisher Scientific). The plasmid DNA was then chemically transformed into specific *E. coli* strains for further work (32). Expression from the pCA24N vector is driven from a T5-*lac* chimeric promoter. In the case of membrane protein expression (TolC and AcrA) the basal expression from uninduced promoter was used in complementation experiments to avoid toxicity of membrane protein overexpression due to the Sec system saturation, whereas expression of Hmp (a cytosolic NO-detoxifying protein) was induced by 1 mM IPTG.

**Table 1:**
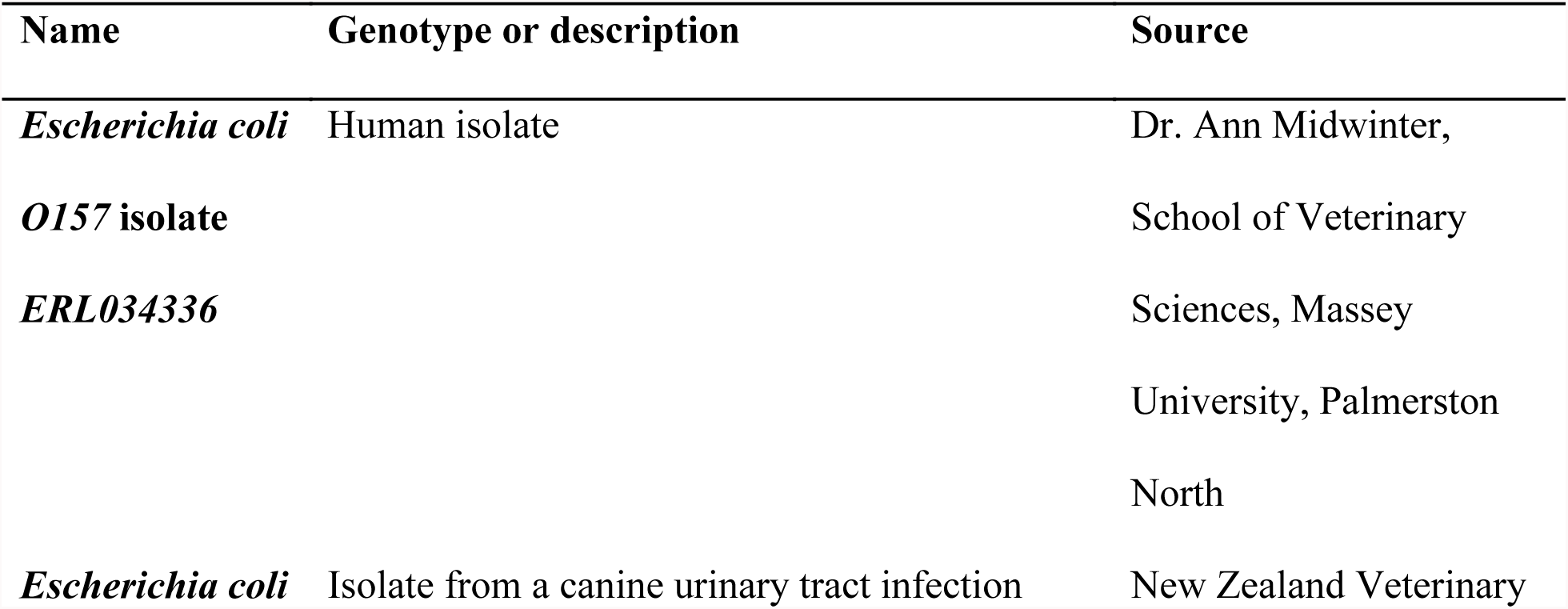
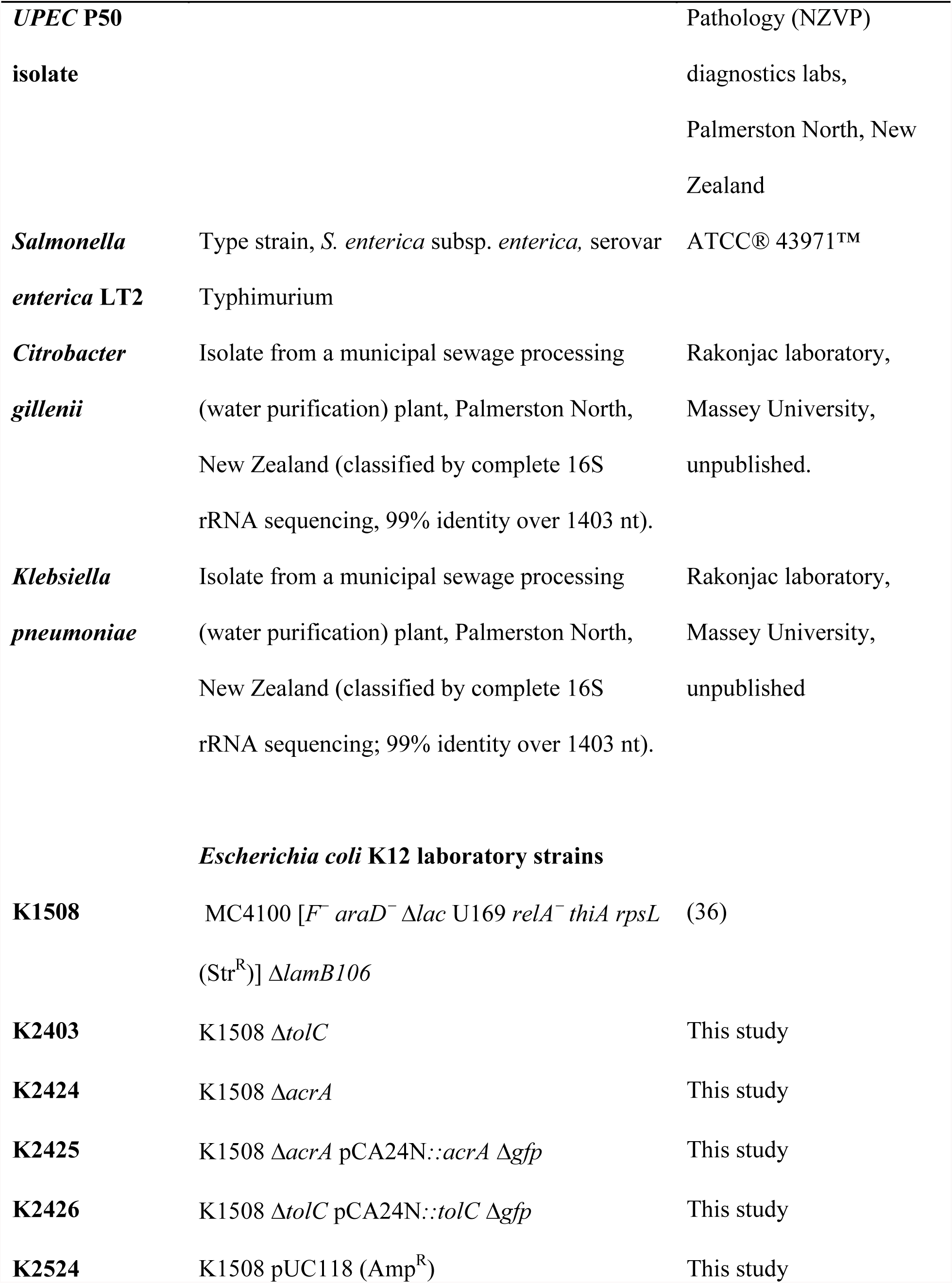
Bacterial strains used in this study

**Table 2:**
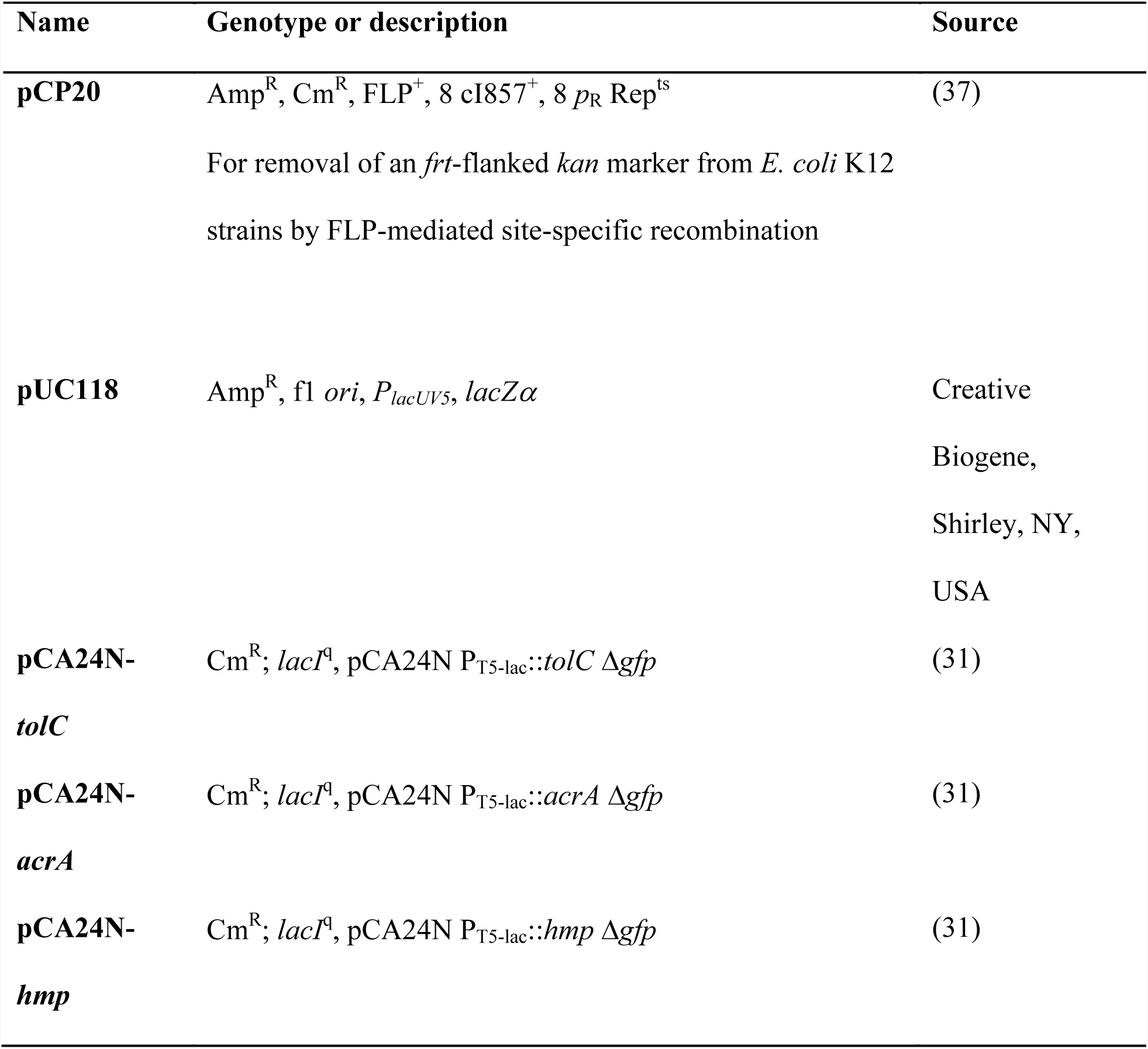
List of plasmids used in this study

Bacterial culture was grown in 2xYT medium (BD Difco) at 37°C with shaking at 200 rpm. For preparation of exponential phase cells, fresh overnight culture was 100-fold diluted and incubated to reach the OD_600nm_ of about 0.1-0.3. This cell suspension was then diluted to the desirable concentration depending on specific purposes. Sodium Deoxycholate was a kind gift from New Zealand Pharmaceuticals Ltd. Antibiotics used in this study were purchased from GoldBio. CM4 was purchased from Enamine (catalog number Z49681516).

### Checkerboard assay

The checkerboard assay for DOC and FZ was performed on the Corning 384-well microtiter plate with the concentration of DOC ranging from 20000 µg/mL to 0 µg/mL and the concentration of FZ ranging from 10 µg/mL to 0 µg/mL, prepared by 2-fold serial dilution. The concentration range could be adjusted depending on the sensitivity of different bacterial strains and the types of nitrofurans to cover at least 2 × MIC to 0.06 × MIC of each drug.

Each well contained the starting inoculum of approximately 10^6^ CFU/mL, 2 % DMSO and the predefined concentration of each drug in the total volume of 50 µL. The wells containing no drugs and 10 µg/mL tetracycline were used as negative controls and positive controls, respectively. After dispensing the reagents, the plate was pulse centrifuged at 1000 × g to eliminate any bubbles. The plate was then incubated at 30°C and the OD_600nm_ of the sample was monitored for every 1 h within 24h using Multiskan^TM^ GO Microplate Spectrophotometer (Thermo Scientific). Each combination was performed in triplicate. The growth inhibition with the cut-off value of 90 % at the time point 24 h was used to define the MIC of the drug used alone or in combination (33). The fractional inhibitory concentration index (FICI) for the two drugs was calculated as follows:

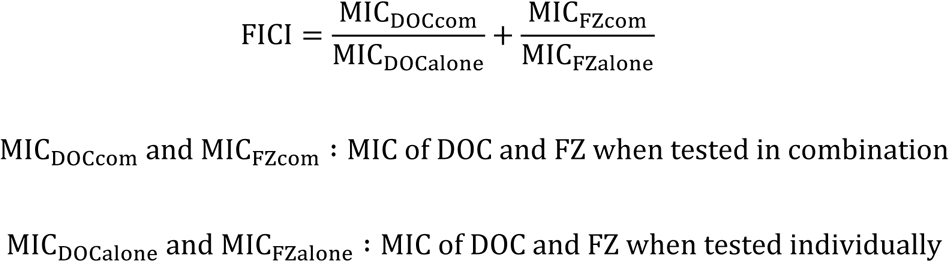

The interaction between two drugs was interpreted as synergistic if FICI was ≤ 0.5, indifferent if it was > 0.5 and ≤ 4, and antagonistic if it was > 4 (34).

### Time kill assay

Exponential phase bacterial culture at about 10^6^ CFU/ml was prepared in the final volume of 10 mL containing 2 % DMSO plus DOC at 2500 µg/ml alone or FZ at 0.5 × MIC µg/mL alone or both drugs. The treatments containing no drug were used as negative controls. The samples were incubated at 30°C with shaking at 200 rpm. At the time points of 0 h, 2 h, 4 h, 6 h, 8 h and 24 h, 500 µL were taken from each treatment and centrifuged at 10000 × g for 15 min before being re-suspended in 100 µL maximum recovery diluent (0.1 % peptone, 0.85 % NaCl). 10 µL of serial dilutions was plated on 2xYT agar followed by overnight incubation at 37°C to determine the cell count. Each treatment was performed in triplicate. The antimicrobial interaction was interpreted as synergistic if the combinatorial treatment caused a killing efficiency ≥2 log higher than the most active agent (35).

## Acknowledgements

We thank Dr. Anne Midwinter, School of Veterinary Sciences, Massey University, for providing a *E. coli* human O157 isolate and New Zealand Veterinary Pathology Ltd. for an isolate of a canine *E. coli* UPEC strain (P50). We are grateful to Fraser Glickman from Rockefeller University High Throughput and Spectroscopy Resource Center for hosting and advice on small-molecule drug screen of a synergy screen and to the National BioResource Project (NBRP) via Genetics Strains Research Center, National Institute of Genetics, Japan, for providing the ASKA collection. The Keio Collection was purchased from Dharmacon *via* ThermoFisher (Australia).

## Funding information

Vuong Van Hung Le has received funding from Callaghan Innovation PhD Scholarship. This work was supported by Massey University, the New Zealand Ministry of Business, Innovation and Employment and New Zealand Pharmaceuticals LTD.

## References

1. O'Neill J, Review-team. 2014. Antimicrobial resistance: tackling a crisis for the health and wealth of nations. https://amr-review.org/,

2. Bollenbach T. 2015. Antimicrobial interactions: mechanisms and implications for drug discovery and resistance evolution. Curr Opin Microbiol 27:1–9.

3. Taneja N, Kaur H. 2016. Insights into newer antimicrobial agents against Gram-negative bacteria. Microbiol Insights 9:9–19.

4. Begley M, Gahan CG, Hill C. 2005. The interaction between bacteria and bile. FEMS Microbiol Rev 29:625–651.

5. Merritt ME, Donaldson JR. 2009. Effect of bile salts on the DNA and membrane integrity of enteric bacteria. J Med Microbiol 58:1533–1541.

6. Cremers CM, Knoefler D, Vitvitsky V, Banerjee R, Jakob U. 2014. Bile salts act as effective protein-unfolding agents and instigators of disulfide stress in vivo. Proc Natl Acad Sci USA 111:E1610–E1619.

7. Nishino K, Yamaguchi A. 2001. Analysis of a complete library of putative drug transporter genes in Escherichia coli. J Bacteriol 183:5803–5812.

8. Van Bambeke F, Glupczynski Y, Plesiat P, Pechere JC, Tulkens PM. 2003. Antibiotic efflux pumps in prokaryotic cells: occurrence, impact on resistance and strategies for the future of antimicrobial therapy. J Antimicrob Chemother 51:1055–1065.

9. Paul S, Alegre KO, Holdsworth SR, Rice M, Brown JA, McVeigh P, Kelly SM, Law CJ. 2014. A single-component multidrug transporter of the major facilitator superfamily is part of a network that protects Escherichia coli from bile salt stress. Mol Microbiol 92:872–884.

10. Sistrunk JR, Nickerson KP, Chanin RB, Rasko DA, Faherty CS. 2016. Survival of the fittest: how bacterial pathogens utilize bile to enhance infection. Clin Microbiol Rev 29:819–836.

11. Chamberlain RE. 1976. Chemotherapeutic properties of prominent nitrofurans. J Antimicrob Chemother 2:325–336.

12. Vass M, Hruska K, Franek M. 2008. Nitrofuran antibiotics: a review on the application, prohibition and residual analysis. Vet Med-Czech 53:469–500.

13. Whiteway J, Koziarz P, Veall J, Sandhu N, Kumar P, Hoecher B, Lambert IB. 1998. Oxygen-insensitive nitroreductases: analysis of the roles of nfsA and nfsB in development of resistance to 5-nitrofuran derivatives in Escherichia coli. J Bacteriol 180:5529–5539.

14. Sandegren L, Lindqvist A, Kahlmeter G, Andersson DI. 2008. Nitrofurantoin resistance mechanism and fitness cost in Escherichia coli. J Antimicrob Chemother 62:495–503.

15. McCalla DR. 1979. Nitrofurans, p 176-213. In Hahn FE (ed), Mechanism of Action of Antibacterial Agents doi:10.1007/978-3-642-46403-4. Springer Berlin Heidelberg, Heidelberg, Germany.

16. McOsker CC, Fitzpatrick PM. 1994. Nitrofurantoin: mechanism of action and implications for resistance development in common uropathogens. J Antimicrob Chemother 33 Suppl A:23-30.

17. Bertenyi KK, Lambert IB. 1996. The mutational specificity of furazolidone in the lacI gene of Escherichia coli. Mutat Res 357:199–208.

18. Roldan MD, Perez-Reinado E, Castillo F, Moreno-Vivian C. 2008. Reduction of polynitroaromatic compounds: the bacterial nitroreductases. FEMS Microbiol Rev 32:474–500.

19. Ona KR, Courcelle CT, Courcelle J. 2009. Nucleotide excision repair is a predominant mechanism for processing nitrofurazone-induced DNA damage in Escherichia coli. J Bacteriol 191:4959–4965.

20. Kumar M, Adhikari S, Hurdle JG. 2014. Action of nitroheterocyclic drugs against Clostridium difficile. Int J Antimicrob Agents 44:314–319.

21. Vumma R, Bang CS, Kruse R, Johansson K, Persson K. 2016. Antibacterial effects of nitric oxide on uropathogenic Escherichia coli during bladder epithelial cell colonization--a comparison with nitrofurantoin. J Antibiot (Tokyo) 69:183–186.

22. Forrester MT, Foster MW. 2012. Protection from nitrosative stress: a central role for microbial flavohemoglobin. Free Radic Biol Med 52:1620–1633.

23. McCollister BD, Hoffman M, Husain M, Vazquez-Torres A. 2011. Nitric oxide protects bacteria from aminoglycosides by blocking the energy-dependent phases of drug uptake. Antimicrob Agents Chemother 55:2189–2196.

24. Martinez-Puchol S, Gomes C, Pons MJ, Ruiz-Roldan L, Torrents de la Pena A, Ochoa TJ, Ruiz J. 2015. Development and analysis of furazolidone-resistant Escherichia coli mutants. APMIS 123:676–681.

25. Darkoh C, Lichtenberger LM, Ajami N, Dial EJ, Jiang ZD, DuPont HL. 2010. Bile acids improve the antimicrobial effect of rifaximin. Antimicrob Agents Chemother 54:3618–3624.

26. Anes J, McCusker MP, Fanning S, Martins M. 2015. The ins and outs of RND efflux pumps in Escherichia coli. Front Microbiol 6:587.

27. Granik VG, Grigoriev NB. 2011. Exogenous nitric oxide donors in the series of C-nitro compounds. Russ Chem Rev 80:171–186.

28. Baba T, Ara T, Hasegawa M, Takai Y, Okumura Y, Baba M, Datsenko KA, Tomita M, Wanner BL, Mori H. 2006. Construction of Escherichia coli K-12 in-frame, single-gene knockout mutants: the Keio collection. Mol Syst Biol 2:2006 0008.

29. Thomason LC, Costantino N, Court DL. 2007. E. coli genome manipulation by P1 transduction. Curr Protoc Mol Biol Chapter 1:Unit 1 17.

30. Datsenko KA, Wanner BL. 2000. One-step inactivation of chromosomal genes in Escherichia coli K-12 using PCR products. Proc Natl Acad Sci U S A 97:6640–6645.

31. Kitagawa M, Ara T, Arifuzzaman M, Ioka-Nakamichi T, Inamoto E, Toyonaga H, Mori H. 2005. Complete set of ORF clones of Escherichia coli ASKA library (a complete set of E. coli K-12 ORF archive): unique resources for biological research. DNA Res 12:291–299.

32. Green R, Rogers EJ. 2013. Transformation of chemically competent E. coli. Methods Enzymol 529:329–336.

33. Campbell J. 2010. High-throughput assessment of bacterial growth inhibition by optical density measurements. Curr Protoc Chem Biol 2:195–208.

34. Odds FC. 2003. Synergy, antagonism, and what the chequerboard puts between them. J Antimicrob Chemother 52:1–1.

35. Doern CD. 2014. When does 2 plus 2 equal 5? A review of antimicrobial synergy testing. J Clin Microbiol 52:4124–4128.

36. Spagnuolo J, Opalka N, Wen WX, Gagic D, Chabaud E, Bellini P, Bennett MD, Norris GE, Darst SA, Russel M, Rakonjac J. 2010. Identification of the gate regions in the primary structure of the secretin pIV. Mol Microbiol 76:133–150.

37. Cherepanov PP, Wackernagel W. 1995. Gene disruption in Escherichia coli: TcR and KmR cassettes with the option of Flp-catalyzed excision of the antibiotic-resistance determinant. Gene 158:9–14.

